# Quantitative PCR (qPCR) vs culture-dependent detection to assess honey contamination by *Paenibacillus larvae*

**DOI:** 10.1101/741801

**Authors:** Simone Crudele, Luciano Ricchiuti, Addolorato Ruberto, Franca Rossi

## Abstract

The detection/quantification in honey of spores of the bacterial pathogen *Paenibacillus larvae*, the etiological agent of the American Foulbrood (AFB) infectious disease of honey bee, represents a useful diagnostic tool to identify apiaries at risk of infection and carry out focused prevention measures. Plate count is currently the recommended method used to analyze presence and number of spores for this pathogen. However, its validity needs to be re-assessed since *P. larvae* field strains have a low rate of germination in culture media.

Therefore, in this study, culture-dependent *P. larvae* detection/quantification was compared with quantitative PCR (qPCR) based analysis for 139 honey samples collected in 2017 and 2018 from as many apiaries in the Abruzzo region. According to qPCR based detection, 58.27% samples were positive for *P. larvae*, while only 33.8 % samples were positive by plate count.

Moreover, the levels of contamination with the two methods differed for most samples. Potential false positives were obtained by plate count for 12.9% samples that were negative with the qPCR test. On the other hand, 26.6% samples were culture negative but qPCR positive, suggesting that not all the field strains were able to develop in plate. This was confirmed by obtaining *P. larvae* growth from those samples after supplementing the medium with germination stimulants.

Results strongly suggested the necessity to improve culture methods or replace them with molecular detection/quantification for a more reliable AFB risk estimation.

## Introduction

American foulbrood (AFB), an infectious disease caused by the bacterial spore former *Paenibacillus larvae*, represents a hard challenge for honey bee survival, as it can destroy bee colonies by killing infected larvae and pupae. It is unsuccessfully contrasted by antibiotic treatments, that do not kill the spores of the etiologic agent thus not preventing their accumulation and persistence in hives for extremely long periods (Alippi et al., 2004). In the European countries antibiotic use for AFB treatment is banned and the spread of *P. larvae* spores is prevented exclusively by burning infected hives and equipment after manifestation of the disease symptoms, a tardive, thus inefficient, measure (Locke et al., 2019). Moreover, a study in the Abruzzo region suggested that the latter practice, though being mandatory by law, is largely disregarded (Ricchiuti et al., 2019). Examination of hive matrices such as honey (Alippi et al., 2004), hive debris (Bassi et al., 2018) and adult bees (Forsgren and Laugen, 2014) for the presence of *P. larvae* spores can allow early diagnosis before clinical manifestation and prevents *P. larvae* massive multiplication and transmission (OIE, 2019).

In particular, the examination of honey has been frequently used to estimate AFB prevalence (Ricchiuti et al., 2019). It was suggested that honey contamination reflects the infection status of bee colonies during the period in which nectar is collected, while in other periods false negative results may be obtained from clinically diseased colonies (Nordström et al., 2002). Nevertheless, honey has particular importance as a diagnostic matrix since it is commercialized individually by many beekeepers, thus representing the most easily available matrix for microbiological testing for AFB risk at beekeeper level and allowing focused inspections for clinical signs in case of high contamination levels. Moreover, contaminated honey is responsible for disease spread at the international level (Alippi, 2004), so that its control for *P. larvae* contamination has relevance for product commercialization.

Currently, the best described method for honey examination for *P. larvae* presence is plate count (OIE, 2019), though it was shown that the germination capacity of *P. larvae* field strains in culture media is poor (Forsgreen, 2008). Quantitative PCR (qPCR) was mentioned as a useful tool to monitor the presence of *P. larvae* in hive matrices. However, no reports on its potentialities in the definition of honey contamination levels are available (OIE, 2019). Therefore, this study was carried out to better evaluate the reliability of culture based methods and qPCR based detection/quantification in providing a faithful insight of the *P. larvae* honey contamination. To this purpose 139 honey samples collected in 2017 and 2018 from as many beekeepers operating in the Abruzzo region were analyzed by both techniques. Presumptive false negatives obtained by plate count were re-analyzed by supplementing the culture medium with germination stimulants according to Alvarado et al., 2013.

## Materials and methods

### Bacterial strains and culture conditions

*P. larvae* ATCC 9545 was used as a positive control in molecular detection/identification tests and to test the fertility of the culture media in each analysis session. *P. larvae* strains were subcultured by streaking on Blood Agar (BA) and incubating at 37°C in the presence of 9.8% CO_2_.

Honey samples were provided by beekeepers conferring commercial samples to the Istituto Zooprofilattico product quality competition, and each sample was from a different beekeeper. Samples were stored at −20°C before analysis.

*P. larvae* counts were carried out twice for each sample as follows: 1 g honey was weighted in a 2 ml sterile microtube and 1 ml sterile deionized water was added; honey was allowed to melt by incubation at 55°C for 5 – 15 min; the honey suspension was incubated at 80°C for 10 min to kill *P. larvae* vegetative forms; the suspension was centrifuged at 10,000 rpm for 5 min; the supernatant was eliminated by aspiration, leaving about 100 µl of it for pellet resuspension; the concentrated spore suspension was spread on a plate of the *P. larvae* agar (PLA) medium described by Schuch et al., 2001. By this procedure, the lowest detectable number of spores was 1 CFU/g of honey.

Honey samples originating more than 300 colonies on the plate were analyzed again by plating the 1:10 dilution of the concentrated suspension.

Colonies of all the different morphologies obtained for each sample and resembling those described for *P. larvae* (Ricchiuti et al., 2019), were isolated by streaking on Blood Agar (BA) plates.

To stimulate germination of *P. larvae* for samples negative to plate count but positive to qPCR, PLA medium was supplemented with 1.2 mM L-tyrosine and 0.2 mM uric acid as germination stimulants according to Alvarado et al., 2013. This was done by spreading 200 µl of a ten-fold concentrated suspension of the two substances in PBS buffer on the surface of PLA plates paying attention to uniformly suspend the insoluble L-tyrosine particles.

### DNA extraction from honey

DNA extraction from honey was carry out by a procedure similar to that described for plate count with the difference that the pellet obtained by centrifugation after thermal treatment at 80°C for 10 min was washed twice with 2 ml of sterile phosphate buffered saline (PBS, 8.0 g/L NaCl, 0.2 g/L KH_2_PO_4_, 2.9 g/L Na_2_HPO_4_, 0.2 g/L KCl, pH 7.4) and then extracted with the NucleoSpin® Tissue extraction kit (Carlo Erba, Milan, Italy) according to the instructions.

Extraction of DNA from single colonies was done as described by Rossi et al. (2018). Crude DNA extracts were 1:100 diluted before use in qPCR.

### qPCR testing

The qPCR method described by Rossi et al. (2018) was used both for direct analysis of honey and for colony identification of presumptive *P. larvae* isolates.

## Results and discussion

The PLA medium, conceived by Schuch et al. (2001), allows the recovery of *P. larvae* in an extent not significantly different from a non-selective medium and therefore was chosen for carrying out *P. larvae* counts in this study.

In Figure 1 the percentages of samples comprised in different count ranges according to plate count and qPCR are shown. It can be noticed that qPCR, opposite to plate count, has not the disadvantage of needing confirmation by identifying the isolates. Moreover, qPCR was more sensitive than plate count, being able to identify more samples with low contamination levels.

**Figure 1.**
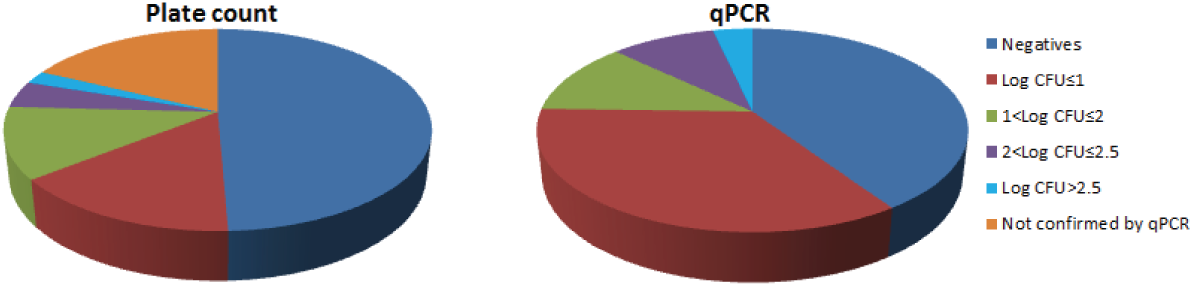
Percentages of honey samples per contamination level according to plate count (a) and qPCR (b).

The superiority of qPCR in reliably identifying honey samples contaminated by *P. larvae* was evident at the single sample level, as presented in Figure 2. Plate count data are reported only for the samples for which isolate identification was confirmed by qPCR.

**Figure 2.**
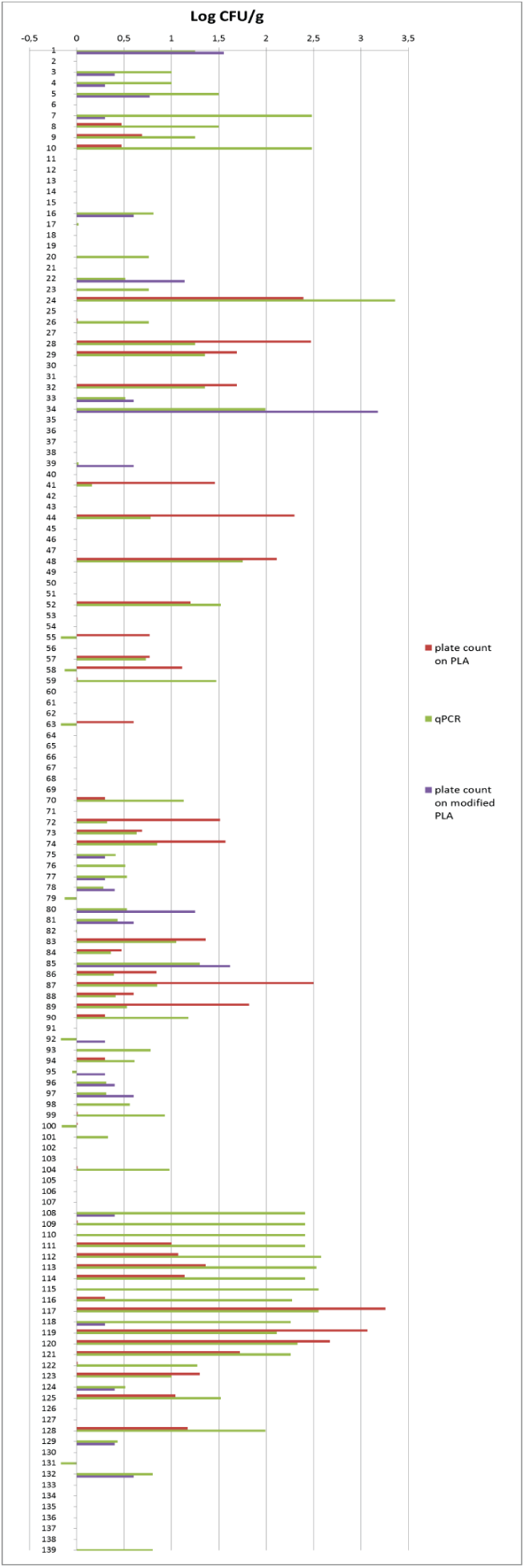
*P. larvae* contamination levels of single honey samples determined by plate count and qPCR. Averages of two measurements are shown.

It can be noticed that all samples positive in cultivation based detection were also identified by qPCR and that additional samples, representing a 26.6% percentage of the total sample number, were positive for *P. larvae* only with the qPCR test. In order to investigate if those samples represented false negatives of the plate count method, a cultivation medium conceived to increase germination capacity of *P. larvae* spores, adapted from a study carried out in different culture conditions, was used to re-analyze those samples. The medium consisted in PLA supplemented with L-tyrosine and uric acid, two substances that had been proven to induce germination of *P. larvae* spores among other compounds found in the bee larva intestine (Alvarado et al., 2013). The modified PLA medium allowed the growth of *P. larvae* in two thirds of the previously culture negative and qPCR positive samples, so that colonies could be isolated and identified by qPCR also from those samples. This indicated that routine use of the modified isolation medium would increase the recovery of *P. larvae*, improve isolation of different strains and provide a more realistic scenario of the contamination status. By using this method the percentage of culture-positive samples raised from 33.8% to 51%.

The inability of culture media not supplemented with germination stimulants to allow growth of all strains could explain why, as found by Bassi et al. (2018) negative microbiological test results can be obtained for honey samples from colonies with AFB symptoms.

The differences in contamination levels found for each sample with the two techniques were probably due to the not complete specificity of the culture medium in the case of samples with counts higher than those determined by qPCR, or to the defective germination of some *P. larvae* strains in the case of samples with counts lower than those determined by qPCR.

In comparison to a previous study carried out three years earlier in the same geographical area (Ricchiuti et al., 2019) by using non-supplemented PLA medium, the percentage of positive samples was similar. Detection/enumeration by qPCR indicated an even higher prevalence, thus showing that not much was done to contrast AFB spread in a region where apiculture is largely practiced (Ricchiuti et al., 2019). This situation is particularly serious, considering that even for low levels of honey contamination AFB clinical signs can occur (Bassi et al., 2018).

It can be concluded that the microbiological diagnosis methods for AFB must be improved for a realistic evaluation of *P. larvae* prevalence and that the application of qPCR, that gives rapidly more reliable results than cultivation procedures, could be preliminarily applied to identify beekeepers with high contamination levels and samples on which to carry out isolations for epidemiological studies.

## References

Alvarado I, Phui A, Elekonich MM, Abel-Santos E. 2013. Requirements for *in vitro* germination of *Paenibacillus larvae* spores. J. Bacteriol. 195:1005–1011. doi: 10.1128/JB.01958-12.

Forsgren E, Stevanovic J, Fries I. 2008. Variability in germination and in temperature and storage resistance among *Paenibacillus larvae* genotypes. Vet. Microbiol. 129:342–349. doi: 10.1016/j.vetmic.2007.12.001.

Locke B, Low M, Forsgren E. 2019. An integrated management strategy to prevent outbreaks and eliminate infection pressure of American foulbrood disease in a commercial beekeeping operation. Prev. Vet. Med. 167:48–52. doi: 10.1016/j.prevetmed.2019.03.023.

Nordström S, Forsgren E, Fries I. 2002. Comparative diagnosis of American foulbrood using samples of adult honey bees and honey. J. Apicult. Sci. 46:5–12.

Ricchiuti L, Rossi F, Del Matto I, Iannitto G, Del Riccio AL, Petrone D, Ruberto G, Cersini A, Di Domenico M, Cammà C. 2019. A study in the Abruzzo region on the presence of *Paenibacillus larvae* spores in honeys indicated underestimation of American foulbrood prevalence in Italy. J. Apicult. Res. 58:416–419. doi.org/10.1080/00218839.2018.1541651

Rossi F, Amadoro C, Ruberto A, Ricchiuti L. 2018. Evaluation of quantitative PCR (qPCR) *Paenibacillus larvae* targeted assays and definition of optimal conditions for its detection/quantification in honey and hive debris. Insects. 9: pii: E165. doi: 10.3390/insects9040165.

Schuch DMT, Madden RH, Sattler A. 2001. An improved method for the detection and presumptive identification of *Paenibacillus larvae* subsp. *larvae spores* in honey. J. Apic. Res. 40, 59–64.

World Assembly of Delegates of the OIE. American Foulbrood of Honey Bees. Manual of Diagnostic Tests and Vaccines for Terrestrial Animals 2019; Chapter 3.2.2. pp. 1–17. Available online: https://www.oie.int/fileadmin/Home/eng/Health_standards/tahm/3.02.02_AMERICAN_FOULBROOD.pdf (accessed on 19 August 2019).

